# Evaluating computational approaches for comparison of protein expression across cancer indications

**DOI:** 10.1101/2024.08.26.609731

**Authors:** Jixin Wang, Xiaowen Tian, Wen Yu, Ben Pullman, John Bullen, Elaine Hurt, Wenyan Zhong

**Author notes:** Jixin Wang and Xiaowen Tian are co-first authors. **Corresponding author:** Wenyan Zhong. AstraZeneca, New York, NY, USA 10036.

## Abstract

**Background:** The National Cancer Institute’s Clinical Proteomic Tumor Analysis Consortium (CPTAC) recently generated harmonized genomic, transcriptomic, proteomic, and clinical data for over 1,000 tumors across 10 cohorts to facilitate pan-cancer discovery research. However, protein expression comparison across CPTAC cohorts remains challenging due to non-uniform missing data and varying protein expression distribution patterns across tumor types. Here, we present our efforts to evaluate various missing data handling and normalization strategies to create a normalized pan-cancer protein expression dataset.

**Results:** First, we developed a novel algorithm to select robustly expressed proteins in tumors within any CPTAC cohort. Second, we applied a cohort hybrid imputation approach to protein abundance values from FragPipe within each cohort based on protein expression distribution patterns. Third, we calculated intensity-based absolute quantification using protein abundance values and applied both global and smooth quantile normalization methods. Our results indicate that global quantile normalization ensured identical distribution across cohorts for both tumor and normal tissues, while smooth quantile normalization preserved distribution differences between biological conditions. We assessed our method by comparing differential protein expression analysis results with and without normalization. Additionally, we examined the ranks of protein expression in the normalized CPTAC dataset for selected proteins with high protein-to-RNA expression correlation across CPTAC cohorts. We then compared these protein expression ranks with their RNA expression ranks across corresponding cohorts in The Cancer Genome Atlas (TCGA). Differential protein expression analysis revealed a high level of agreement in the fold change of tumor versus normal tissue within cohorts before and after normalization. Furthermore, our results indicate that global quantile normalization resulted in the highest cohort rank correlation between CPTAC and TCGA for selected proteins.

**Conclusions:** In summary, our thorough analysis demonstrates that global quantile normalization surpasses both smooth quantile normalization and no normalization, as evidenced by its higher rank correlation across cancer cohorts between CPTAC and TCGA for selected proteins. The findings suggest that combining cohort hybrid imputation with global quantile normalization is an effective method for creating a normalized CPTAC pan-cancer protein dataset, which can facilitate the study of protein expression across different cancer types.

## BACKGROUND

The identification and prioritization of therapeutic targets are pivotal in both drug development and biological understanding. Traditionally, target prioritization has been predominantly tailored to specific cohorts. However, recent developments in oncology underscore the significance of indication prioritization, where a single target holds relevance across multiple cancer types, thereby streamlining drug development and expediting the creation of novel therapies.

The Clinical Proteomic Tumor Analysis Consortium (CPTAC), established by the National Cancer Institute [1], stands as a comprehensive resource of global mass-spectrometry based proteomics experiments. These initiatives enable high-throughput biological understanding by elucidating the relative and absolute expression of nearly the entire proteome across over 10 tumor types, inclusive of both tumor tissue and normal adjacent tissue (NAT). The data generated by CPTAC has proven to be instrumental in target discovery and prioritization within tissue cohorts, leveraging a methodology to estimate absolute protein abundance from relative quantitation data [2–4].

However, a substantial challenge in cross-indication selection lies in comparing protein expression levels across CPTAC cohorts due to a diverse array of samples, each exhibiting distinct missing data and expression patterns. Considering this, we present our endeavors to assess and validate various methodologies for handling missing data and normalization, with the aim of producing a normalized pan-cancer protein expression dataset. Such a dataset has the potential to serve as a valuable resource for the development of cancer therapeutics.

## METHODS

### Evaluation of CPTAC pan-cancer reprocessed proteomics data

CPTAC pan-cancer samples were reprocessed using the FragPipe computational platform (version 15) with MSFragger (version 3.2) and Philosopher (version 3.4.13) [5] for protein identification and quantitation as well as iBAQ derivation, as described in Wang et al. [3]. FragPipe was used to assess protein abundance in tumor and normal tissues from 10 cancer indications in CPTAC. Log2-transformed protein abundance was used for assessing missing value patterns and developing an imputation strategy. “Cohort” was defined as the unique combination of tissue type and indication, and “missing rate” was defined as the percentage of samples with missing values in each cohort.

### Algorithm for selection of robustly expressed proteins

An algorithm was developed to select robustly expressed proteins in the samples as follows. For protein *j* in tumor samples from indication *i*, the percentage of samples with missing abundance values was denoted as M_{*ij*} and the median abundance from non-missing observations was denoted as A_{*ij*}. The corresponding percentile rank of A_{*ij*} within each cohort *i* was obtained and denoted as F_*i*{(A_{*ij*})}. The protein *j* was kept in all cohorts if there was an indication *i* such that F_*i*{(A_{*ij*})} > 50% and M_{*ij*} < 25%. In other words, the protein was kept if the median abundance percentile rank was >50% and the corresponding missing rate was <25% in at least one indication among tumor samples. The determination of these cutoffs was empirical, aimed at achieving a balance between discarding an excessive number of proteins and retaining those with a high rate of missing protein expression measurements.

### iBAQ conversion and normalization after cohort hybrid imputation

The R package imputeLCMD [6] was used for imputation after filtering proteins that did not pass the above criteria. This package works via a method selection approach that first determines whether the protein was affected by a missing-not-at-random (MNAR) missingness mechanism, based on the assumption that the distributions of the mean values of protein abundance follow a normal distribution within the cohort [18]. If the determined missing mechanism was MNAR, quantile regression imputation of left-censored data was applied. Otherwise, k–nearest neighbor (KNN) imputation was applied. Imputation was performed on robustly expressed proteins as determined by the method described above, in log2 scale. iBAQ values were then calculated by dividing the raw imputed protein abundance by the number of theoretically observable peptides of the protein from FragPipe, as described by Wang et al. [3]. The iBAQ values were then normalized using one of two quantile normalization methods: (1) global quantile normalization with quantile normalization [7] applied to all samples across indications and tissue types together; or (2) smooth quantile normalization [8] with the assumption that the statistical distribution of each sample was the same within indications or tissue types, but allowing for differences between groups. Additionally, riBAQ (relative iBAQ) [9] and riBAQ-derived copy number [10] were calculated according to Wang et al. [3]. The iBAQ values obtained from the imputed protein abundance was used to derive riBAQ and copy number values without applying additional normalization methods.

### Differential protein expression analysis with normalized iBAQ

To determine whether our missing data imputation and normalization strategy affected downstream analyses, we compared the fold change in differentially expressed proteins (DEP) between tumor and matched normal adjacent tissue (NAT) samples for each indication, using non-normalized, global quantile normalized, and smooth quantile normalized protein iBAQ values.

FragPipe protein abundance–derived iBAQ data from CPTAC tumor and NAT samples were used for DEP analysis using R (version 4.1.1), with empirical Bayes statistics on protein-wise linear models with limma [11] embedded in the DEP package (version 1.16.0) [12]. The correlations between the log2 fold change from DEP analysis were examined between each comparison using Pearson correlation.

### Comparison of the indication ranks of selected proteins between CPTAC and TCGA

We identified proteins whose protein and RNA expression were highly correlated across CPTAC cohorts. Within each indication, we calculated the Pearson correlation between protein and RNA expression using tumor tissues. Then, proteins with a correlation greater than 0.5 across the majority of indications were kept for subsequent analysis. We then compared their protein expression rank across CPTAC cohorts with their RNA expression rank across corresponding cohorts from The Cancer Genome Atlas (TCGA) [13]. Specifically, the median log2(iBAQ) before and after normalization, the median log2(riBAQ) of CPTAC and the median log2(TPM) of TCGA were calculated for those proteins within each indication. Next, indications were ranked by the median log2(iBAQ) before and after normalization, the median log2(riBAQ) and the median log2(TPM) of CPTAC and TCGA, respectively. A weighted rank correlation approach [14] was used to measure rank agreement between each comparison, where the correlation between the protein and RNA expression in CPTAC was used as the weight.

## RESULTS

### Evaluation of CPTAC pan-cancer reprocessed proteomics data

We used the FragPipe computational platform to assess protein abundance in tumor tissue and normal tissue samples from 10 cancer indications in CPTAC. Log2-transformed protein abundance was used to identify missing value patterns and develop an imputation strategy.

The workflow used in this study is outlined in Fig. 1. We developed an algorithm to select robustly expressed proteins across cohorts as described in the methods section. A total of 15,762 proteins were identified in the union of all cohorts, with 8,419 commonly detected proteins across all indications. The protein identifications per cohort at 1% protein-level false discovery rate (FDR) are shown in Supplementary Table 1. In addition to the 8,419 proteins identified in all cohorts, we included 1,718 proteins that, while not present in all cohorts, are considered robustly expressed in certain indications by our robustly expressed proteins selection algorithm defined in the methods section. Finally, we selected 10,137 proteins for further analysis and the selected proteins per cohort are shown in Supplementary Table 2. Moreover, when we assess the total iBAQ values for each sample, excluding the filtered proteins versus including them, our algorithm shows a mere 1% reduction (see Supplementary Fig. S1).

**Fig. 1.**
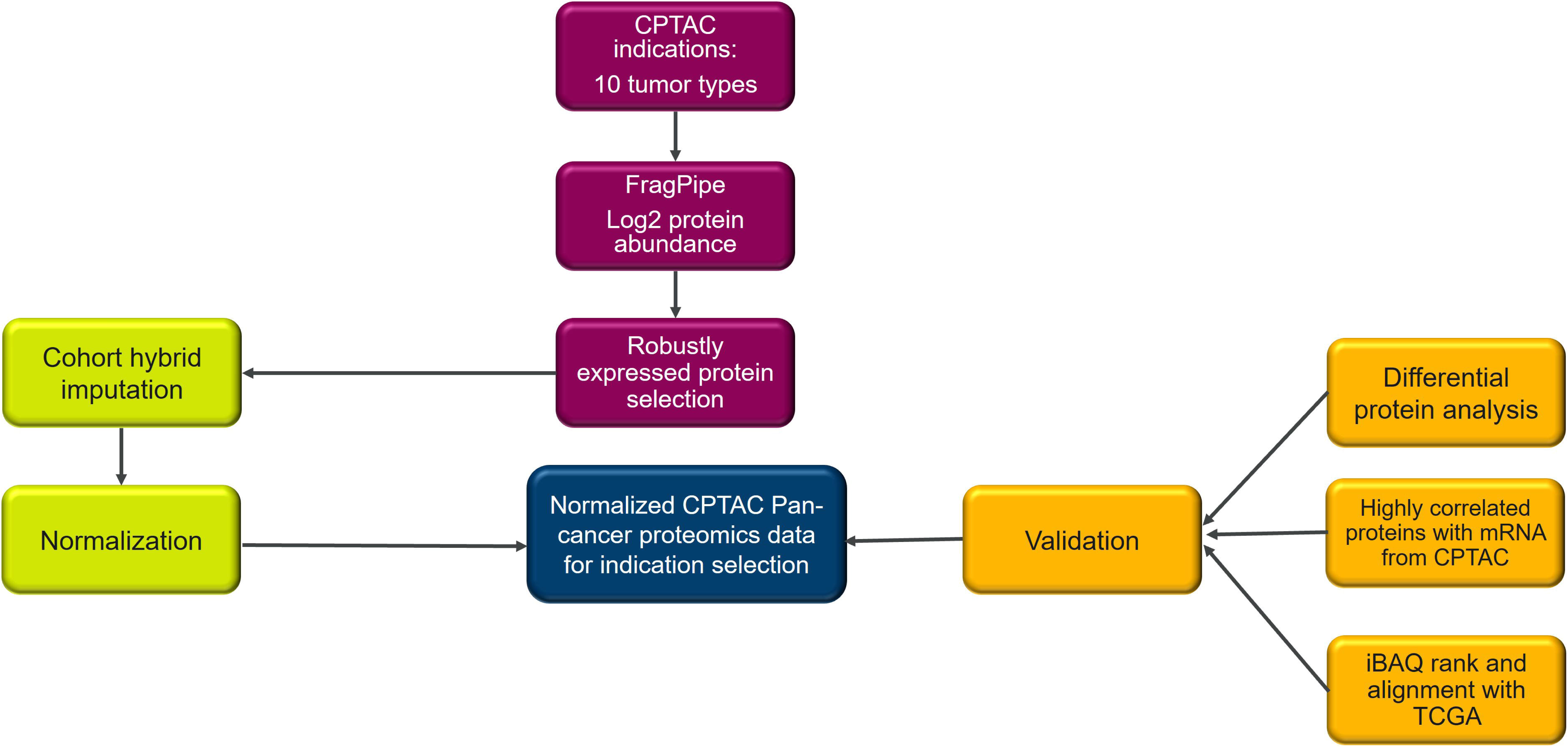
Workflow for a computational approach that enables the comparison of protein expression across CPTAC cohorts. Tissue samples from 10 cancer indications in CPTAC were processed through FragPipe. Protein abundance data were analyzed by iBAQ to derive estimated absolute protein abundance. A cohort hybrid imputation strategy was developed to address missing values. Quantile normalization was applied to iBAQ values. Imputed and normalized iBAQ values were used for validation with differential protein analysis, ranking indications by protein expression and comparing these ranks with those derived from TCGA RNA-Seq data.

### Evaluation of missing values pattern

Our evaluation of the missing values showed both missing-at-random (MAR) and missing-not-at-random (MNAR) patterns within and across cohorts (Fig. 2a and 2b). As shown in Fig. 2a, the COAD cohort had the highest missing values among all cohorts. The most prominent categories of proteins with missing values are those with a missing rate of 0% to 25% in the MAR pattern and those with a missing rate of 75% to 100% in the MNAR pattern. Interestingly, BRCA exhibits a markedly higher number of proteins with the MNAR pattern in normal tissue compared to tumor tissue (Fig. 2b). Furthermore, proteins with higher missing rates tended to have lower expression levels (Supplementary Fig. S2), suggesting an MNAR pattern. Because it was critical to apply the appropriate missing value imputation method before performing any further downstream analysis, we chose a cohort hybrid algorithm to impute missing values [6].

**Fig. 2.**
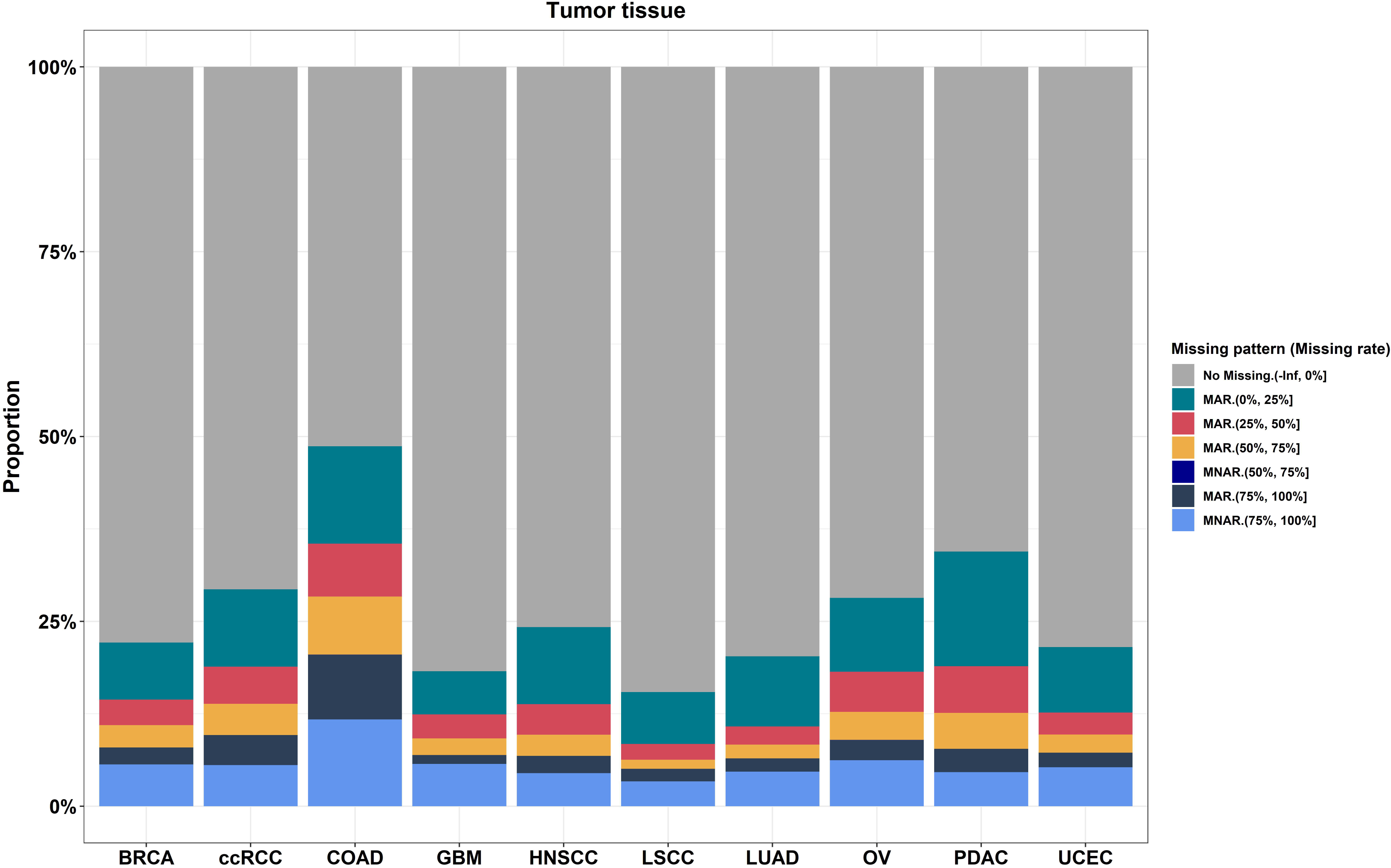
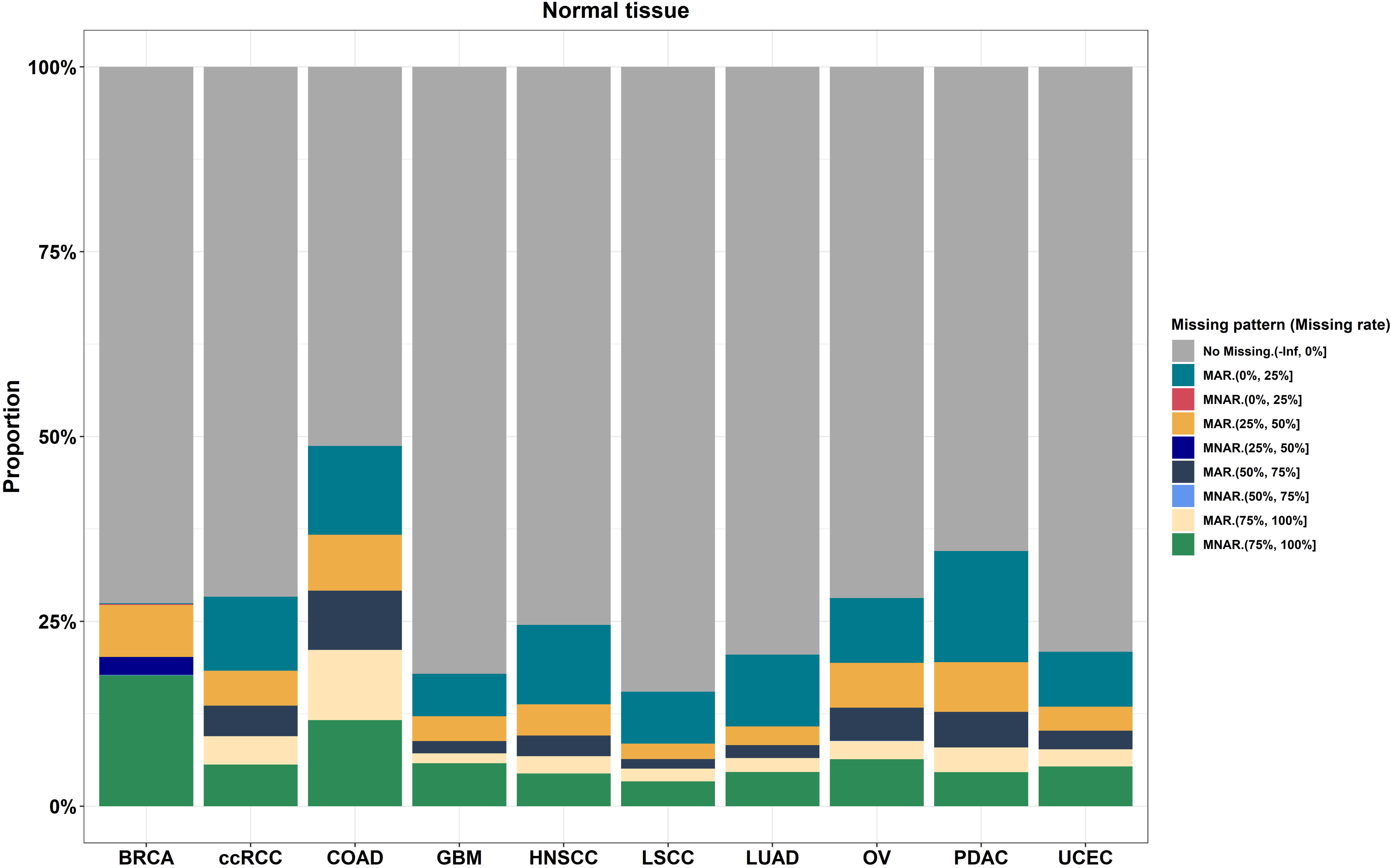
CPTAC missing values pattern assessment in **a** tumor and **b** normal tissue. “Cohort” was defined as the unique combination of tissue type and indication. “Missing rate” was defined as the percentage of samples with missing values in each cohort.

### Assessment of normalization methods

Global quantile normalization ensured identical distribution across cohorts for both tumor and normal tissues, whereas smooth quantile normalization preserved variability across cohorts (Fig. 3a and 3b). The PCA analysis of protein expression across pan-cancer indications revealed that cohort separation based on protein expression is more distinct than tissue types, both before and after normalization. The most noticeable tissue separation occurred following global quantile normalization (Supplementary Fig. S3).

**Fig. 3.**
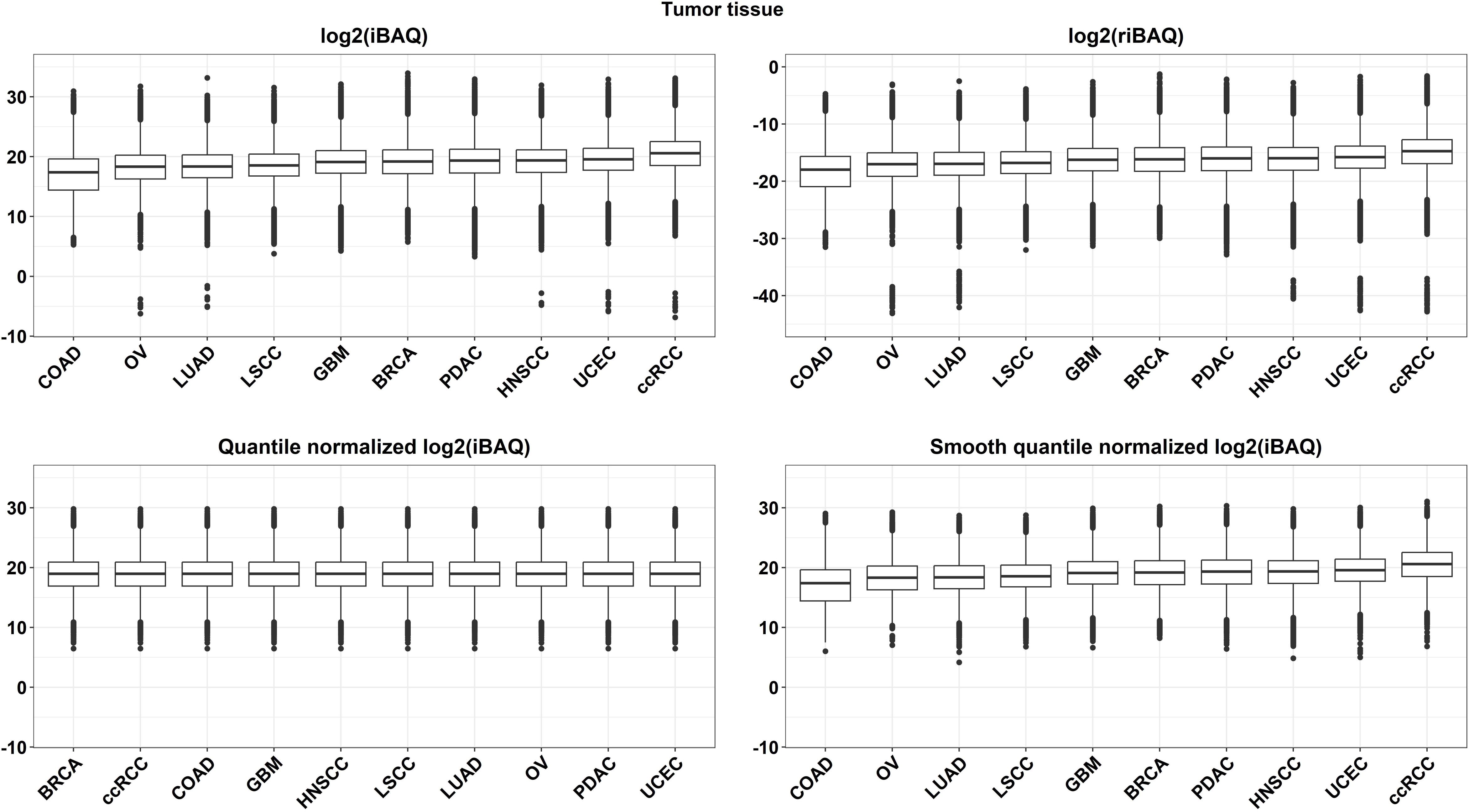
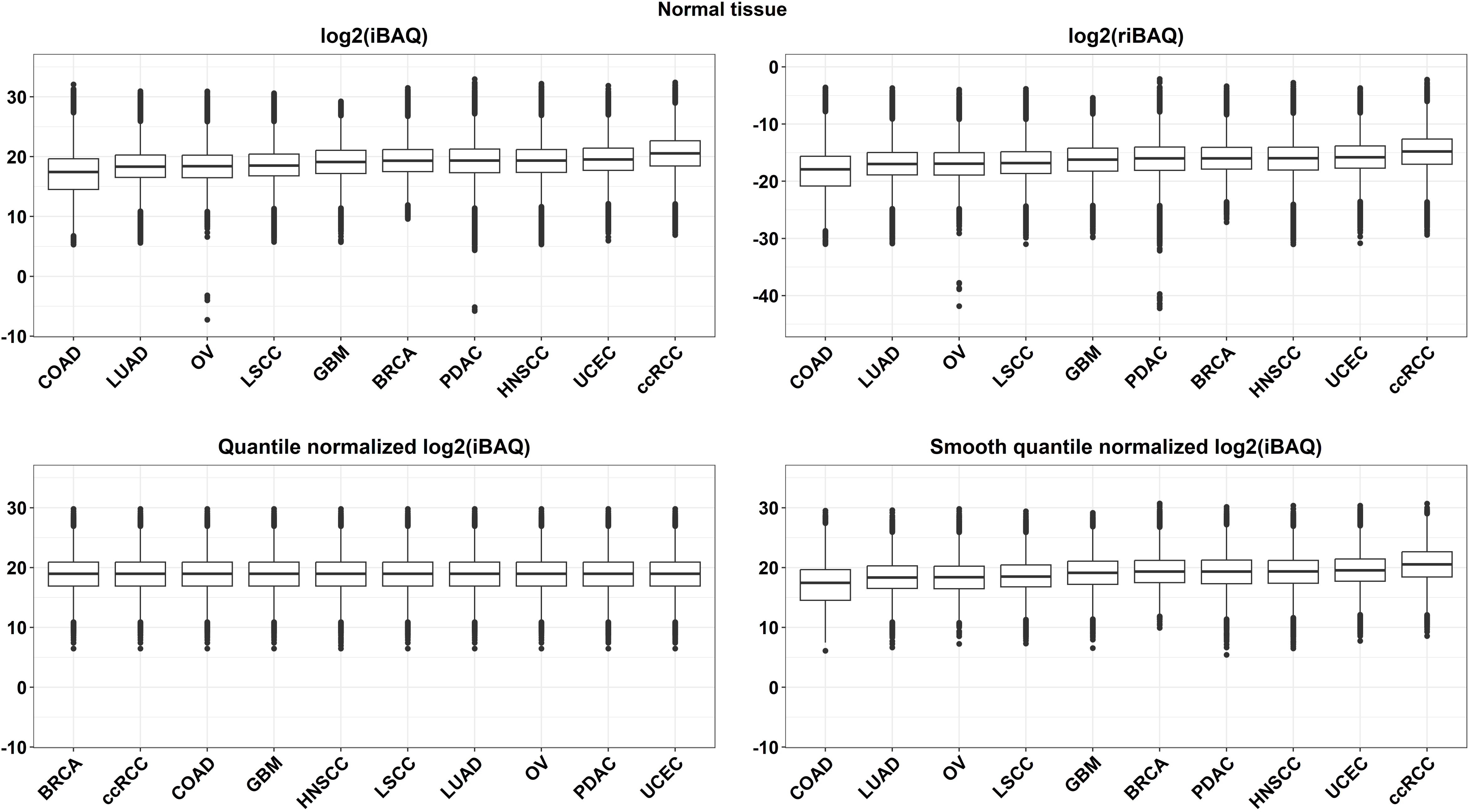
Quantile normalization. **a** Quantile normalization applied to derived iBAQ values after imputation in **a** tumor and **b** normal tissue. Two quantile normalization methods, global and smooth quantile normalization, were used.

### Differential protein expression analysis with normalized and imputed iBAQ values

To determine whether our missing data imputation and normalization strategy affected downstream analyses, we compared the fold change in DEP between tumor tissue and matched normal tissue of selected cohorts (ccRCC, COAD, LSCC, and LUAD), using non-normalized, riBAQ, global quantile normalized, and smooth quantile normalized protein iBAQ values. Our results (Pearson *r* values) demonstrate a strong correlation in fold change between riBAQ normalized data and non-normalized data (ccRCC, *r* = 0.99823; COAD, *r* = 0.99791; LUAD, *r* = 0.99749; LSCC, *r* = 0.99813) (Fig. 4a), global quantile normalized data and non-normalized data (ccRCC, *r* = 0.97262; COAD, *r* = 0.97882; LUAD, *r* = 0.99105; LSCC, *r* = 0.99142) (Fig. 4b). Similar results were observed when we compared smooth quantile normalized data with non-normalized data (ccRCC, *r* = 0.99863; COAD, *r* = 0.99993; LUAD, *r* = 0.99819; LSCC, *r* = 0.99996) (Fig. 4c). The Pearson *r* values tend to decrease as more variability is removed from the dataset; for example, the smooth quantile normalization resulted in *r* = 0.99863 while the global quantile normalized resulted in *r* = 0.97262 in ccRCC. However, with global quantile normalization, the correlations are still greater than 0.97 in the four selected indications, indicating that the global quantile normalization method retained biological differences between tumor tissues and matched normal tissues within cohorts.

**Fig. 4.**
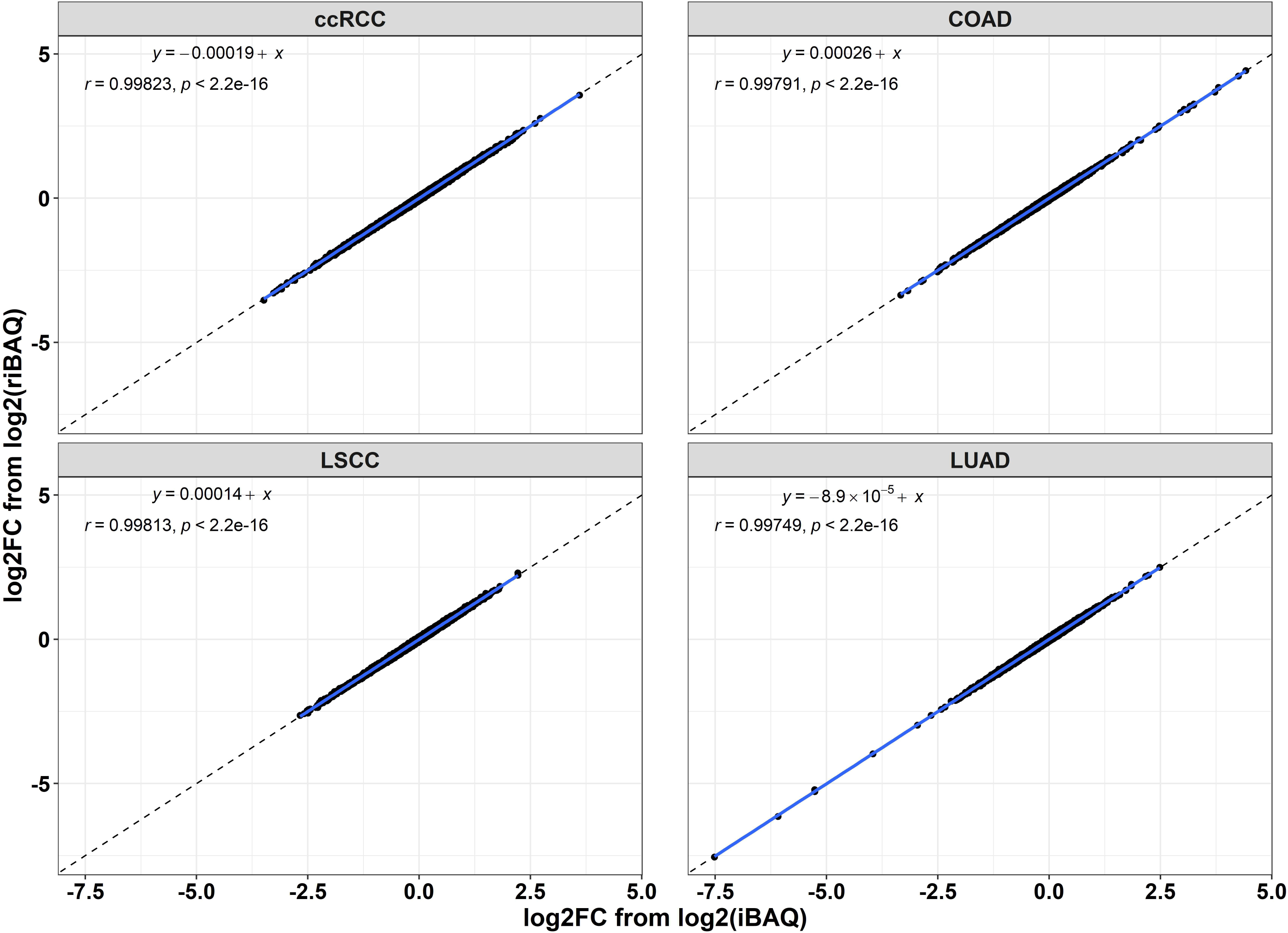
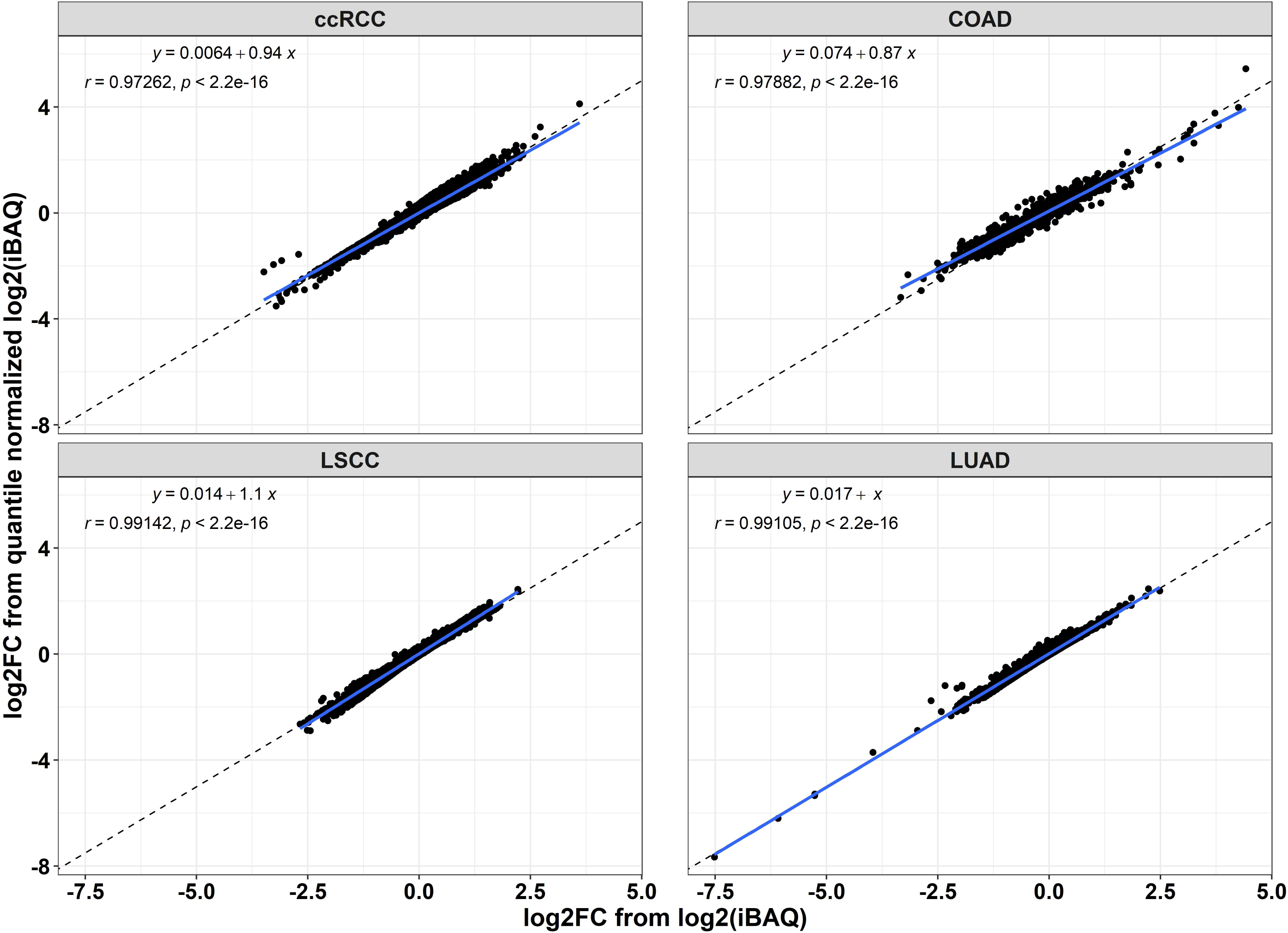
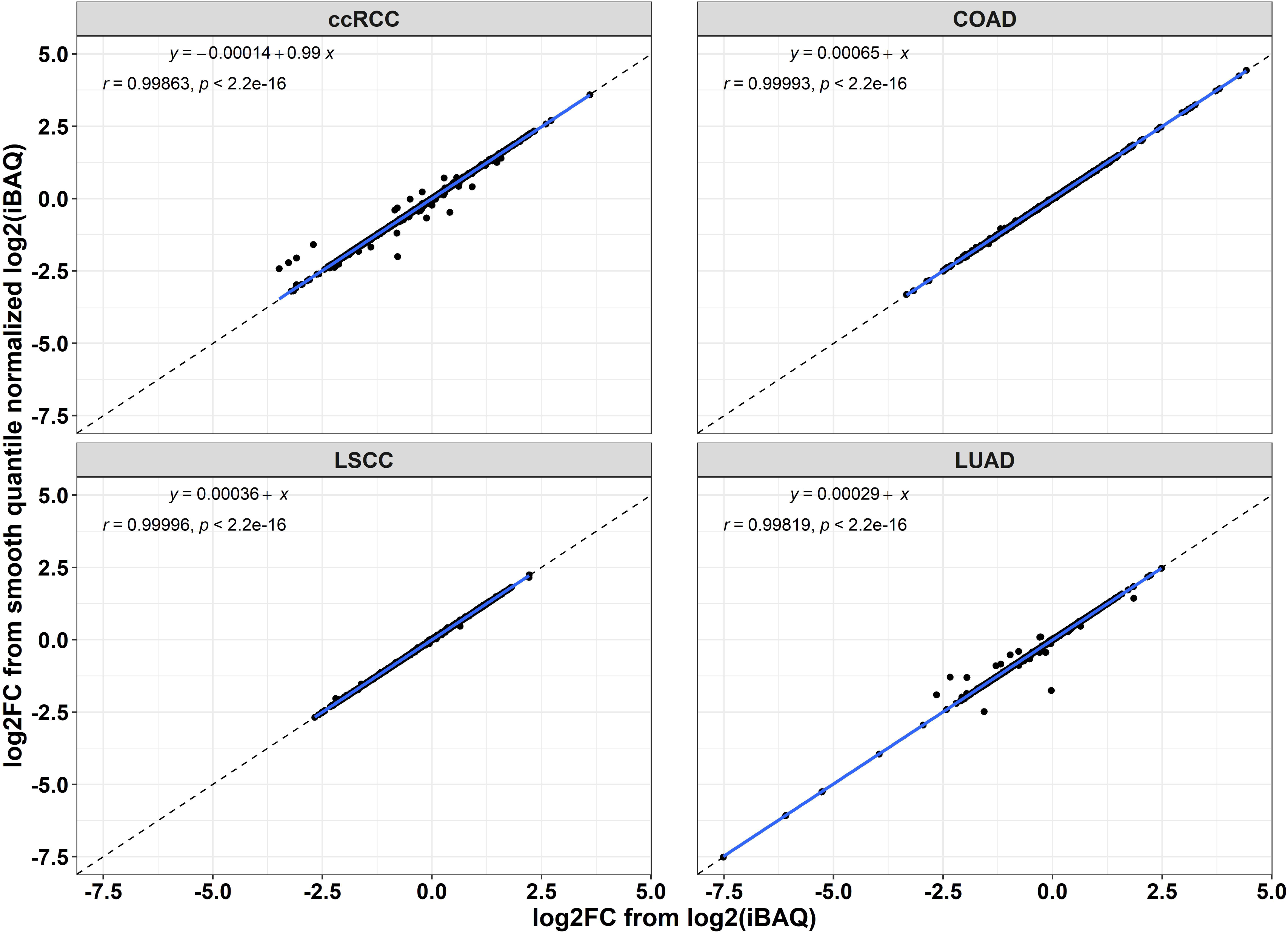
Correlation between fold change from differential expression analysis. **a** riBAQ normalization and no normalization. **b** Global quantile normalization and no normalization. **c** Smooth quantile normalization and no normalization.

### Comparison of the indication ranks of selected proteins between CPTAC and TCGA

We identified proteins whose protein and RNA expression were highly correlated (*r* > 0.5) across CPTAC cohorts: ERAP2, CA9, GSTM3, MX1, and STAT1. We then compared their median protein expression rank across CPTAC cohorts with their median RNA expression rank across corresponding TCGA cohorts. A weighted rank correlation approach [14] was used to measure rank agreement between each comparison (e.g., denoted as “v” in the Supplementary Fig. S4). Ovarian cancer (OV) and glioblastoma multiforme (GBM) are excluded from the weighted rank correlation calculation because they have a small number of proteins with *r* > 0.5, and those proteins do not overlap with those selected from other indications. Global quantile normalization had a higher rank correlation (weighted rank correlation of 0.597–0.931) than smooth quantile normalization (weighted rank correlation of 0.168–0.76) and no normalization (weighted rank correlation of 0.168–0.76) (Supplementary Fig. S4).

## DISCUSSION

CPTAC recently generated harmonized genomic, transcriptomic, proteomic, and clinical data for over 1,000 tumors in 10 cohorts to facilitate pan-cancer discovery research [2]. To use these data for prioritizing cancer surface antigens and selecting indications in the discovery and development of drug targets, protein expression levels often need to be compared and ranked across cohorts [15, 16]. Efforts to do so are hindered, however, by non-uniform missing data and varying protein expression distribution patterns across tumor types. To evaluate various strategies for missing data handling and normalization for the generation of a normalized pan-cancer protein expression dataset, we built a computational workflow that incorporates the selection of robustly expressed proteins, cohort hybrid imputation, and quantile normalization [17]. We then evaluated our strategy by comparing indication ranking between CPTAC and TCGA using a set of proteins with high correlation between protein and RNA expression (Fig. 1). To our knowledge, this work represents the first cross-cohort normalized CPTAC pan-cancer data containing estimated absolute protein expression quantification derived from mass spectrometry–based proteomic data.

There are considerable challenges in harmonizing proteomics data across the 10 CPTAC cohorts since they have been generated over multiple years and analyzed by various labs utilizing different processing pipelines. To avoid potential confounding factors, it was necessary to reanalyze TMT global proteomics data with the same pipeline. We first estimated the absolute protein abundance in reprocessed CPTAC pan-cancer data as described previously [18]. Additionally, to create a unified dataset, it is necessary to define a set of proteins that maximizes the identification of proteins across all 10 cohorts with high quality while minimizing the occurrence of missing values. We designed a protein selection algorithm to effectively balance these requirements. Our robustly expressed proteins selection algorithm defined 10,137 proteins, which indicated that more than half of the human proteome (assuming 20,000 protein products) [19, 20] is robustly expressed.

Proteomics data often contain missing values due to a variety of technical factors, such as protein abundance below the instrument limit of detection, poor ionization efficiency, and low signal-to-noise ratio [21–23]. When merging the 10 cohorts, we faced similar issues. Although many algorithms have been developed for missing value imputation[24], it is critical to select the ones that are best suited to the specific characteristics of the data and downstream analysis needs. An analysis of the missing values in the CPTAC cohort uncovered patterns indicative of both Missing at Random (MAR) and Missing Not at Random (MNAR). Based on these observations, a cohort-hybrid imputation approach was applied. Our study introduces a general framework to manage missing data when comparing protein expression across cohorts. For other specific analyses, one can adopt this model, integrating multiple imputation methods to measure the uncertainty linked to imputation, thereby obtaining more accurate estimations.

Due to the lack of a definitive list of proteins with known expression levels across different tumor types, we chose to use proteins that exhibit a strong correlation between their RNA and protein expression levels. We then assessed our approach by comparing the rank order of these proteins’ expression in our normalized dataset to their RNA expression rank in The Cancer Genome Atlas (TCGA) as an indirect method of evaluation. Furthermore, we have implemented a weighted rank correlation method that accounts for the correlation between protein and RNA expression levels. Through our validation method, we’ve shown a significantly enhanced rank correlation with the application of global cohort normalization as opposed to using a dataset without normalization (see Supplementary Fig. S4). Further, when comparing the fold change of tumor versus matched normal tissues using the normalized dataset versus the original dataset, there was significant agreement (Fig. 4), indicating that our normalization method did not inadvertently eliminate biological variation.

In conclusion, our assessment indicates that a combination of cohort hybrid imputation and global quantile normalization is a reasonable approach to generate a normalized CPTAC pan-cancer protein dataset that could be leveraged to compare protein expression across different cancer types. Nonetheless, the limited number of proteins and indications used for validation in our study represents an important limitation. While our approach was tested using selected highly correlated proteins and rank alignment with TCGA RNA-Seq data, we recommend further validation using a larger number of proteins with expression levels measured across cohorts using other orthogonal experimental methods such as targeted proteomics to enhance the robustness of our approach.

## CONCLUSIONS

We have established a computational workflow for cross-indication comparison from proteomics data, thereby providing a unique data resource to interrogate protein expression across different cancer types. Our results demonstrate that missing data imputation and normalization strategies do not affect downstream analyses. The methodology and the data sources used in this study can serve as valuable resources for cancer research.

## Supplementary Information

The online version contains supplementary material available at[https://doi.org/10.5281/zenodo.13376456].

**Additional file 1. Supplemental Table 1.** The number of proteins and subjects in the pan-cancer indication.

**Additional file 2. Supplemental Table 2.** The number of selected proteins in each cohort.

**Additional file 3. Supplemental Figure 1.** Percentage change in the sum of iBAQ values for each sample.

**Additional file 4. Supplemental Figure 2.** Protein expression missing pattern.

**Additional file 5. Supplemental Figure 3.** Principal component analysis of protein expression across the pan-cancer indication.

**Additional file 6. Supplemental Figure 4.** Comparison of expression ranks of highly correlated proteins in CPTAC and TCGA.

## Acknowledgments

We gratefully acknowledge the CPTAC for providing open-source proteomics data and Deborah Shuman of AstraZeneca for editing the manuscript and formatting the figures. We also acknowledge the presentation of a portion of this work at the AACR Annual Meeting 2024 ( April 5-10, 2024, San Diego; https://aacrjournals.org/cancerres/issue/84/7_Supplement).

## Availability of data and materials

The imputed and normalized iBAQ values of FragPipe output of protein abundance estimation for the 10 CPTAC indications used in this work can be downloaded from https://doi.org/10.5281/zenodo.13376456.

## Authors’ contributions

JW and WZ conceived and designed the study. JW, XT, YW, BP, JB, EH, and WZ analyzed and interpreted the data. All authors contributed to the writing, review, and/or revision of the manuscript, have approved the final version of the manuscript, and agree to be accountable for all aspects of the work.

## DECLARATIONS

### Ethics approval and consent to participate

Not applicable.

### Consent for publication

Not applicable.

## Abbreviations

BRCA: breast cancer
ccRCC: clear-cell renal cell carcinoma
COAD: colon adenocarcinoma
CPTAC: Clinical Proteomic Tumor Analysis Consortium
DEP: differential protein analysis
GBM: glioblastoma multiforme
HNSCC: head and neck squamous-cell carcinoma
iBAQ: intensity-based absolute quantification
riBAQ: relative intensity-based absolute quantification
KNN: k–nearest neighbor
LSCC: lung squamous-cell carcinoma
LUAD: lung adenocarcinoma
MNAR: missing not at random
NAT: normal adjacent tissue
OV: ovarian cancer
PDAC: pancreatic ductal adenocarcinoma
QRILC: quantile regression imputation of left-censored data
TMT: tandem mass tag
UCEC: uterine corpus endometrial carcinoma

