## Supplementary Information for "Evaluating computational approaches for comparison of protein expression across cancer indications"

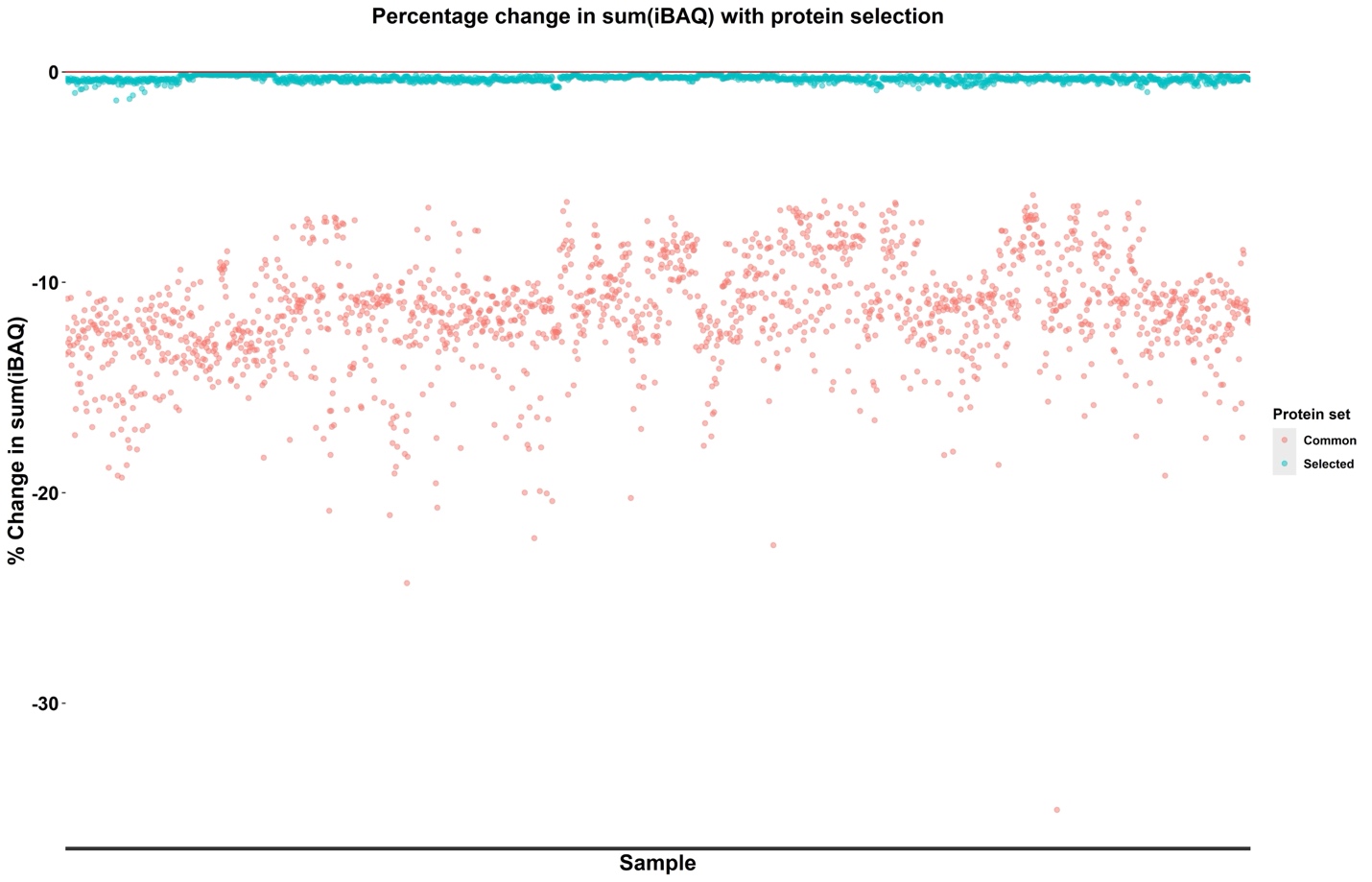

**Fig. S1**	Percentage change in the sum of iBAQ values for each sample. The robustly selected proteins are denoted by “Selected” and the proteins commonly detected across all indications are denoted by “Common” in the figure.

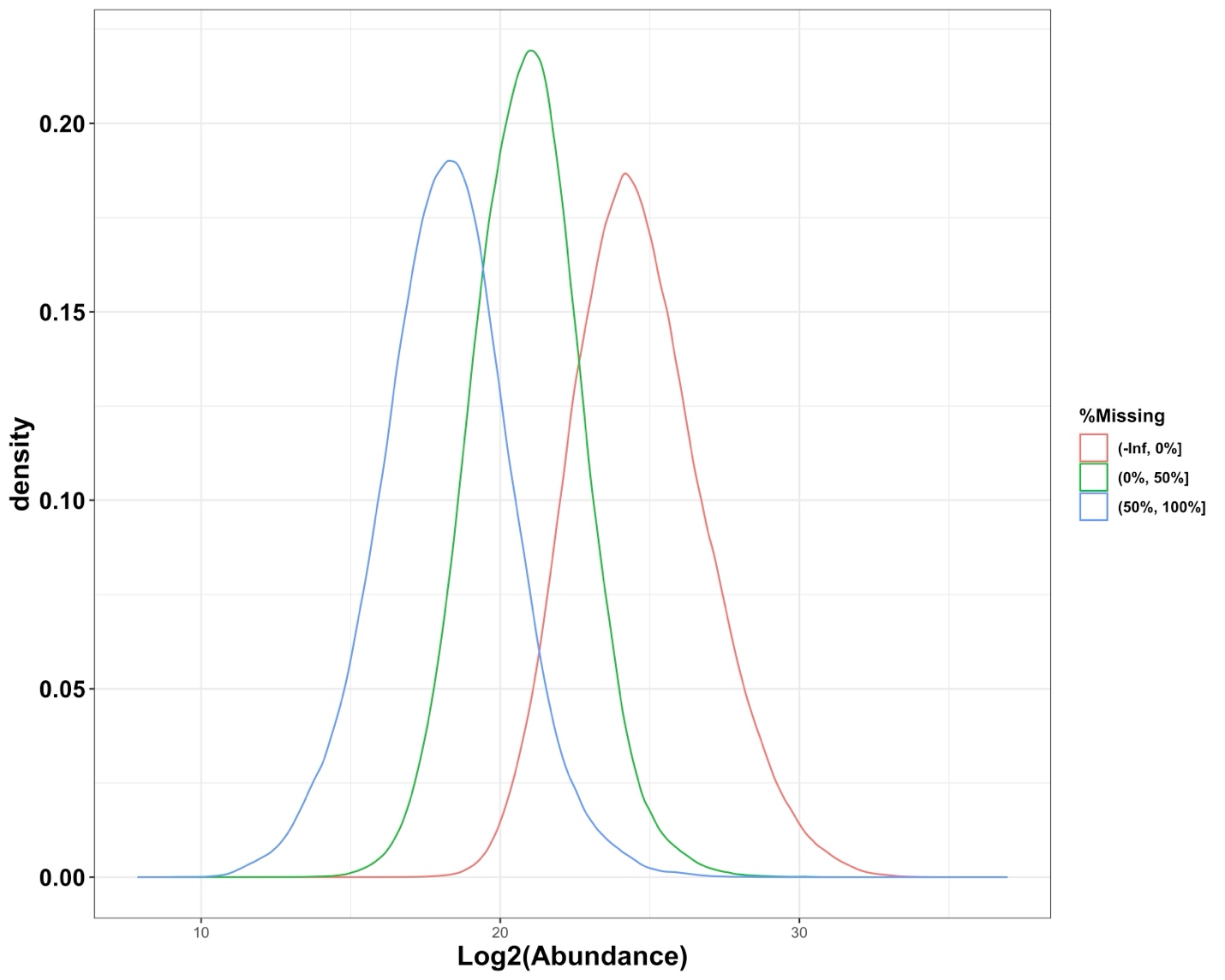

**Fig. S2**	Protein expression missing pattern in the National Cancer Institute’s Clinical Proteomic Tumor Analysis Consortium (CPTAC) dataset.

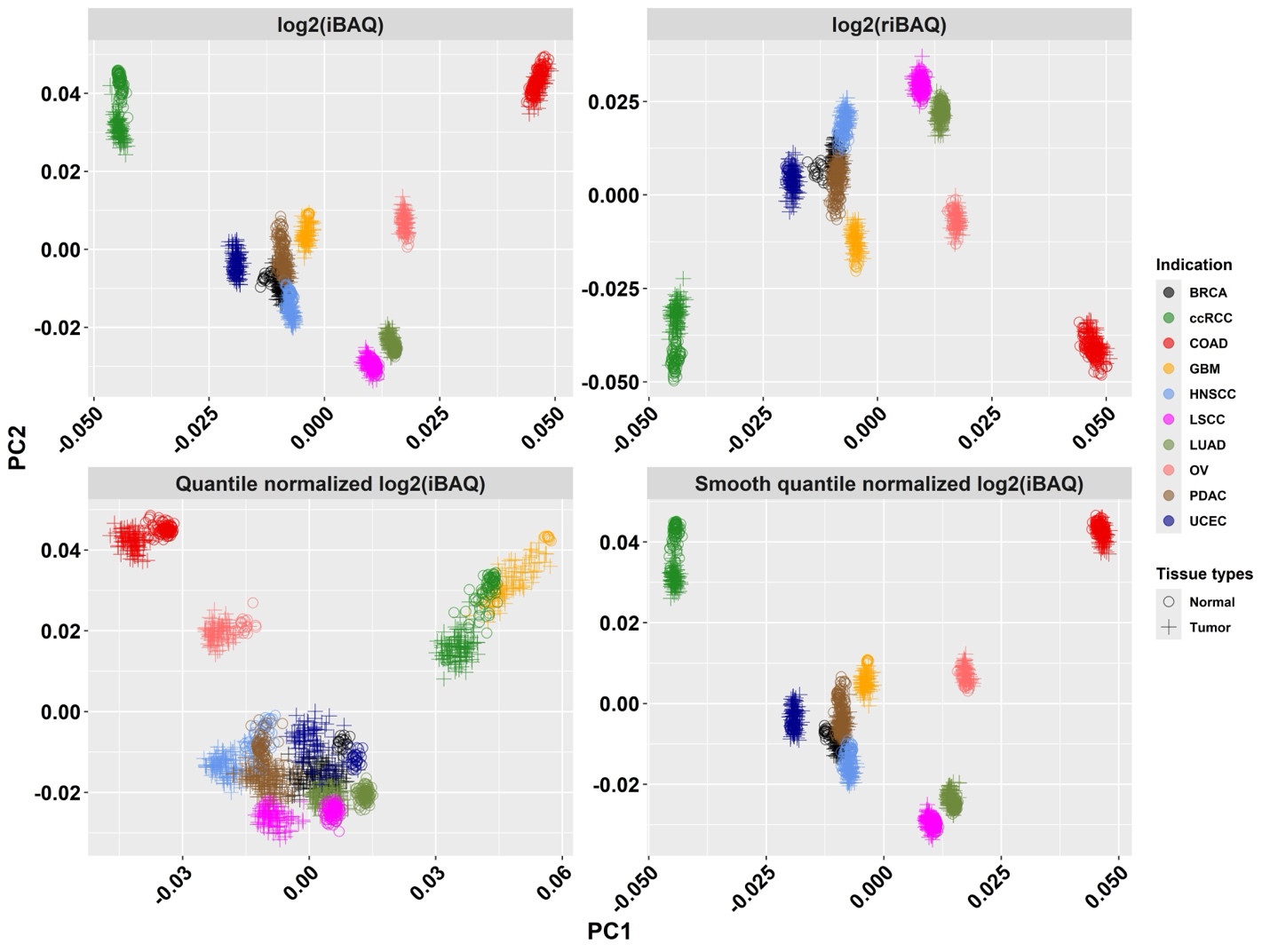

**Fig. S3**	Principal component analysis of protein expression across pan-cancer indications in the National Cancer Institute’s Clinical Proteomic Tumor Analysis Consortium (CPTAC) dataset. Data points are color coded by indication and shaped according to tissue type.

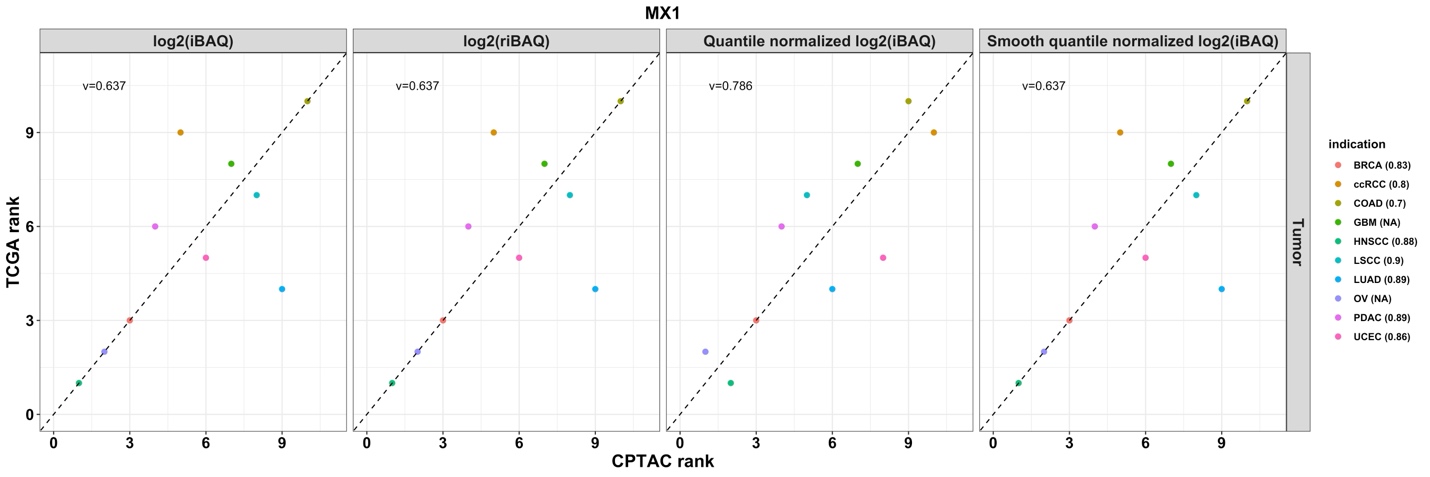

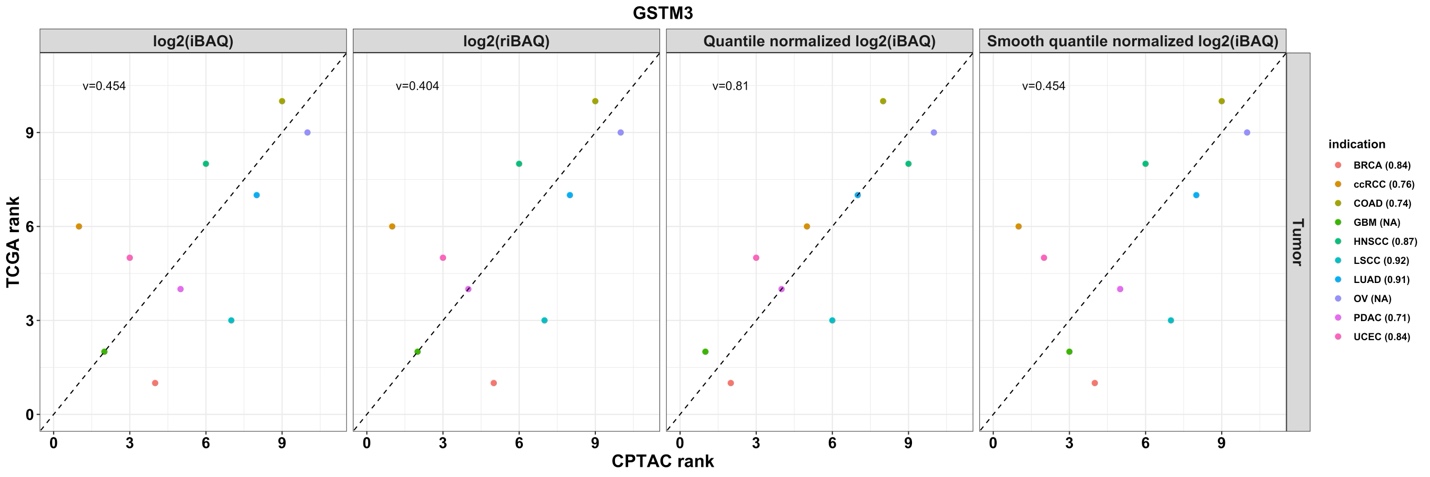

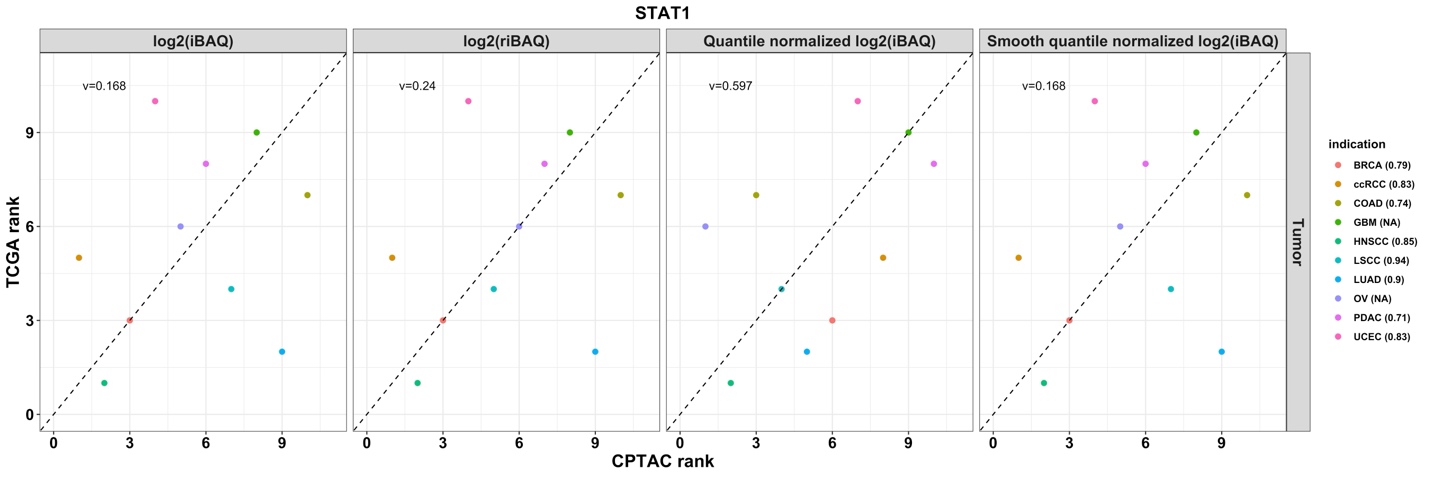

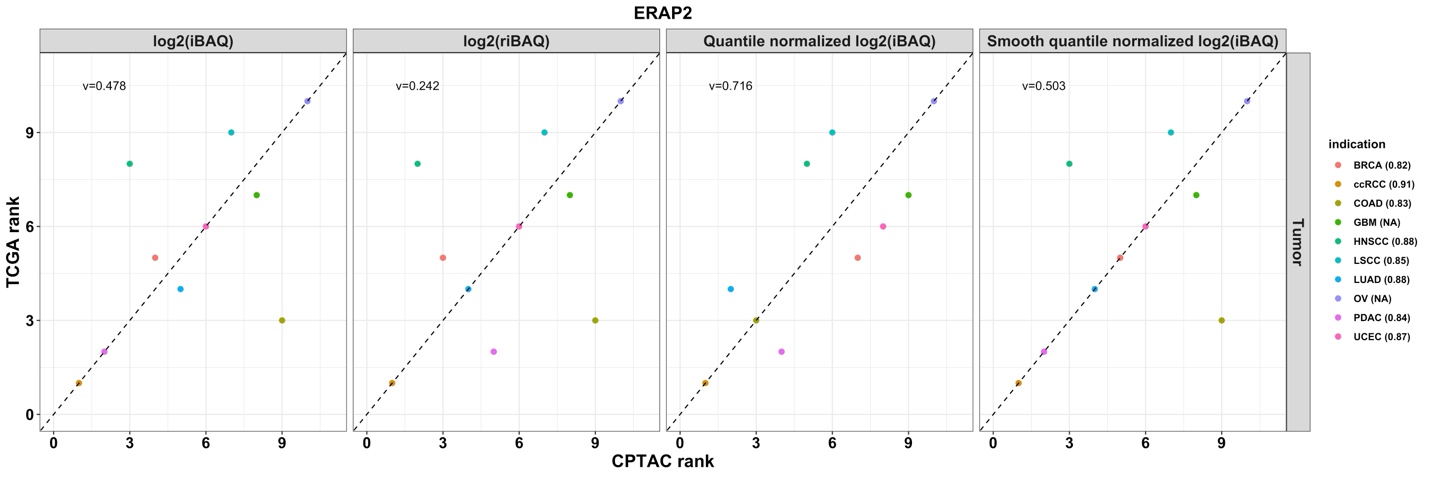

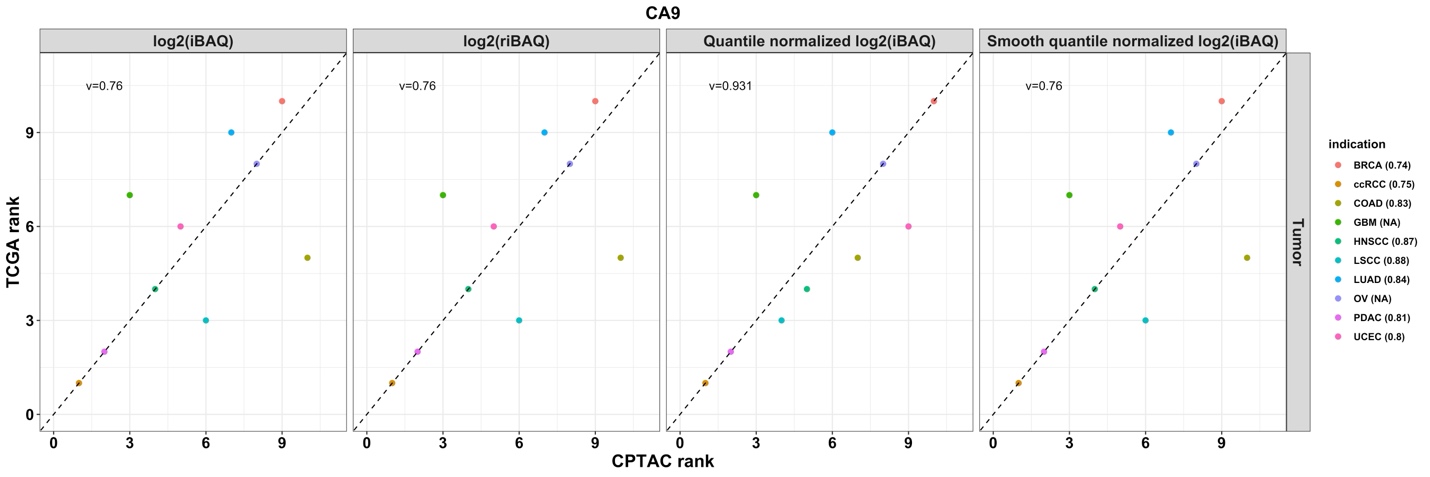

**Fig. S4**	Comparison of expression ranks of highly correlated proteins in CPTAC and TCGA. Data points are color coded by indication and the number in parentheses is the Pearson correlation coefficient between protein and RNA expression in CPTAC. v: the weighted rank correlation coefficient. Protein and RNA expression ranks from Ovarian cancer (OV) and glioblastoma multiforme (GBM) were plotted but excluded from the rank correlation calculation due to the limited number of proteins showing high correlation with RNA, as well as the absence of overlap with proteins selected from other indications.
